# *In situ* profiling of bacterial growth activity during the early stages of *Pseudomonas aeruginosa* infection in airway epithelia

**DOI:** 10.1101/2025.11.13.688211

**Authors:** Albert Fuglsang-Madsen, Janus Anders Juul Haagensen, Claudia Antonella Colque, Helle Krogh Johansen, Søren Molin

## Abstract

*Pseudomonas aeruginosa* is an opportunistic pathogen of major clinical importance, which frequently gives rise to persistent antibiotic resilient infections. To investigate the connection between the bacterial physiological activity and the host environment during the early stages of bacterial infections, we have established a dual-fluorescent reporter system to monitor growth activity of *P. aeruginosa* during infection of a human airway epithelial model. This approach enables quantitative and spatially resolved analysis of bacterial growth within distinct infection micro-niches. Using this infection model, we compared the infection routes for the reference strain PAO1 as well as for two patho-adaptive mutants, PAO1*ΔpscC* (*ΔpscC*) and PAO1*ΔmexZ* (*ΔmexZ*). All three strains colonized apical-, intracellular-, interepithelial-, and epithelial barrier breach sites, but with strain-specific patterns of localisation. PAO1 was rarely observed intracellularly, *ΔpscC* was detected in few cases at epithelial-barrier breach- and interepithelial sites, and *ΔmexZ* colonized apical and intracellular niches in only few cases. Measurements of bacterial growth activities in these niches further revealed distinct hierarchies of bacterial growth activity among the tissue sites: PAO1 bacteria were most active at epithelial-barrier breach sites, *ΔpscC* at interepithelial sites, and *ΔmexZ* in intracellular niches.

Taken together, these data support a model in which *P. aeruginosa* follows a progressive infection continuum from apical colonization to barrier breach, involving interepithelial spread, whereas the mutant strains represented truncated or altered versions of this infection program. More broadly, this study demonstrates the utility of unstable fluorescent reporters for capturing dynamic, niche-specific growth activity patterns during host-pathogen interactions, with implications for both basic pathogenesis research and therapeutic development.

**Author Summary:** In this study, we set out to better understand how bacterial infections develop and progress on human tissues. To do so, we created fluorescent “reporter” strains of *Pseudomonas aeruginosa*, a bacterium that commonly infects the lungs of people with cystic fibrosis and other chronic lung diseases. These genetically modified bacteria allow us to directly observe their growth activity in living human cell cultures using advanced microscopy. Here, we can see how the bacteria grow and spread at different locations within the tissue, and if there are any sites they prefer relative to others. We can also see if those patterns change, depending on any mutations the bacteria might have. In the future, the same genetic engineering can be applied to other bacteria, and the fluorescence can inform us of a range of important bacterial functions; for instance, at precisely what point of the infection process – and where in the tissues – the bacteria produce toxins, are stressed, produce antibiotic-resistance proteins or divide, for example.

Our experiments revealed that some genetic mutations, that are often found in bacterial isolates from hospital infections, change the preference of colonization to distinct tissue sites and confer specialized and unique patterns – and peaks – of growth activity. Some bacterial strains tend to grow on the surface of the tissue, while others are more likely to move between or inside human cells. These behaviours reflect how bacteria evolve during long-term infections and adapt to different environments within the body tissues. The approach we present here provides a new tool for studying how bacterial infections unfold and may ultimately help identify more effective ways to treat or control them.

## 1. Introduction

In the quest to combat infectious diseases and antimicrobial resistance development, researchers need a powerful and translative tool to visualize the dynamic progression of infection by bacterial pathogens. With this in mind, we developed reporter strains, using *Pseudomonas aeruginosa* as a model organism, with the goal to apply them on *in vitro* human cell cultures, set to mimic relevant conditions of the human airways.

Understanding the microscopic dynamics of bacterial infections in *in vitro* human cell cultures is critical for elucidating the mechanisms that underlie pathogenic behaviour (1–5). Such insights enable the generation of hypotheses on why different bacterial species and strains exhibit distinct infection pathways. These spatio-temporal behaviours may be linked to, for instance, nutrient availability, surface adhesion preferences, and/or susceptibility to antibiotics. While transcriptomic and proteomic approaches provide valuable information on gene expression and protein abundance, they often lack the spatio-temporal resolution needed to observe *in situ* gene regulation, as it occurs. In contrast, the use of fluorescent reporter strains offers a powerful non-destructive alternative, allowing real-time *in situ* visualization of promoter activity and thus transcriptional- or mechanistic responses to environmental changes (6). Reporter strains can reveal dynamic expression patterns of genes involved in metabolism and growth (7), virulence factor production (e.g., adhesins, toxins), biofilm development (8–11), or motility (12); crucial traits that influence infection outcomes. By integrating such reporter systems into microscopy-based platforms with cultured human tissue, researchers gain the ability, not only to monitor infection progression, but also to test the effects, and in some cases even elucidate mechanisms of small molecules or other therapeutic agents in real time (13,14). Such therapeutic agents can be evaluated for their capacity to inhibit pathogenic behaviours like biofilm formation or virulence expression, or to target and kill both actively growing- and dormant bacterial subpopulations. Ultimately, the development of such platforms holds great promise for advancing our molecular understanding of bacterial infections and for accelerating the discovery of improved antimicrobial strategies.

In this study, we introduce a dual-reporter system into the standard laboratory strain *P. aeruginosa* PAO1; a representative “naïve” environmental isolate similar to isolates that typically establish initial airway infections in people with cystic fibrosis (pwCF) or chronic obstructive pulmonary disease, before acquiring host-adaptive mutations. Alongside PAO1, we employed two PAO1-derived strains carrying clinically relevant patho-adaptive mutations: a Δ*mexZ* mutant, the most frequently observed patho-adaptation in early infecting clinical CF isolates, and a Δ*pscC* mutant, exemplifying a common adaptive loss of the type III secretion system (T3SS). Loss of T3SS function represents a particularly relevant adaptation, as mutations in T3SS components or in upstream regulators such as *retS* are frequently observed in clinical CF isolates (15). While *pscC* mutations are associated with strongly reduced virulence, a classical hallmark of patho-adaptation in clinical CF isolates, *mexZ* mutations affect regulation of the MexXY efflux pump, which has been shown to indirectly lead to an overproduction of the lectin, LecA, and a distinct colonization pattern *in vitro* on a human airway epithelial model (16).

Human epithelial tissues comprise diverse micro-niches, each with distinct nutrient availability, physicochemical properties, and mechanical forces, which can lead to heterogenous bacterial subpopulations (17,18). For example, oxygen gradients and shear stresses differ markedly between the epithelium’s apical surface, the intercellular space and the intracellular environment, while successful colonization between cells seem to require additional adhesion factors (16). Even genetically identical bacteria can diversify into phenotypically distinct subpopulations in response to such local conditions, mediated by gene regulatory changes (7). Assuming that PAO1, Δ*pscC*, and Δ*mexZ* can all colonize these micro-niches, an open question is whether their subpopulations behave similarly in each site, or if the patho-adaptive mutations fundamentally reshape their growth strategies.

Because both *ΔpscC* and *ΔmexZ* patho-adaptive mutations are known to give rise to markedly different infection phenotypes, one characterized by attenuated T3SS-dependent virulence and intracellular colonization (19–22), the other by interepithelial colonization, altered efflux pump regulation and accompanying pleiotropic effects, we speculated that they would also differ in their tissue-site preferences for growth. This raised three central questions that guided our study: (i) do these patho-adaptive mutations confer differences in preferred micro-niches for each strain? (ii) do the strains exhibit similar growth activities when colonizing each micro-niche? and (iii) if there are preferred micro-niches, are these the areas where the strains exhibit their peak growth activities?

The dual-reporter system was designed to monitor transcription activity of a ribosomal promoter, providing a proxy for bacterial growth (23–26). By directly comparing PAO1 with its pathoadaptive derivatives, we identified strain-specific micro-niche preferences and obtained eco-physiological insights into infection dynamics at the microscale, using image analysis.

## 2. Results

### 2.1 Differences in tight junction breakage and CFU numbers in tissue compartments after infection by the bacterial strains

To assess how patho-adaptive mutations influence infection dynamics, we first examined epithelial integrity and bacterial distribution in the *in vitro* airway model after infection. Measurements of trans-epithelial electric resistance (TEER) and colony-forming (CFU) counts across tissue compartments provided an overview of strain-specific differences in epithelial disruption and bacterial spread. TEER measurements and CFU numbers revealed differences between the PAO1, Δ*mexZ* and Δ*pscC* reporter strains, as expected (27). For TEER measurements, Δ*pscC* showed a non-significant change after 16 h of incubation, in comparison to the uninfected control (Mock) as reported previously (27–29). In contrast, PAO1 and Δ*mexZ* conferred strongly reduced post-infection TEER (Figure 1A). Concerning the bacterial localization in the tissue, the PAO1 population particularly colonizes the apical and attached compartments compared to the mutant strains, which in contrast are less frequently observed in the basolateral compartment (Figure 1B).

**Figure 1:**
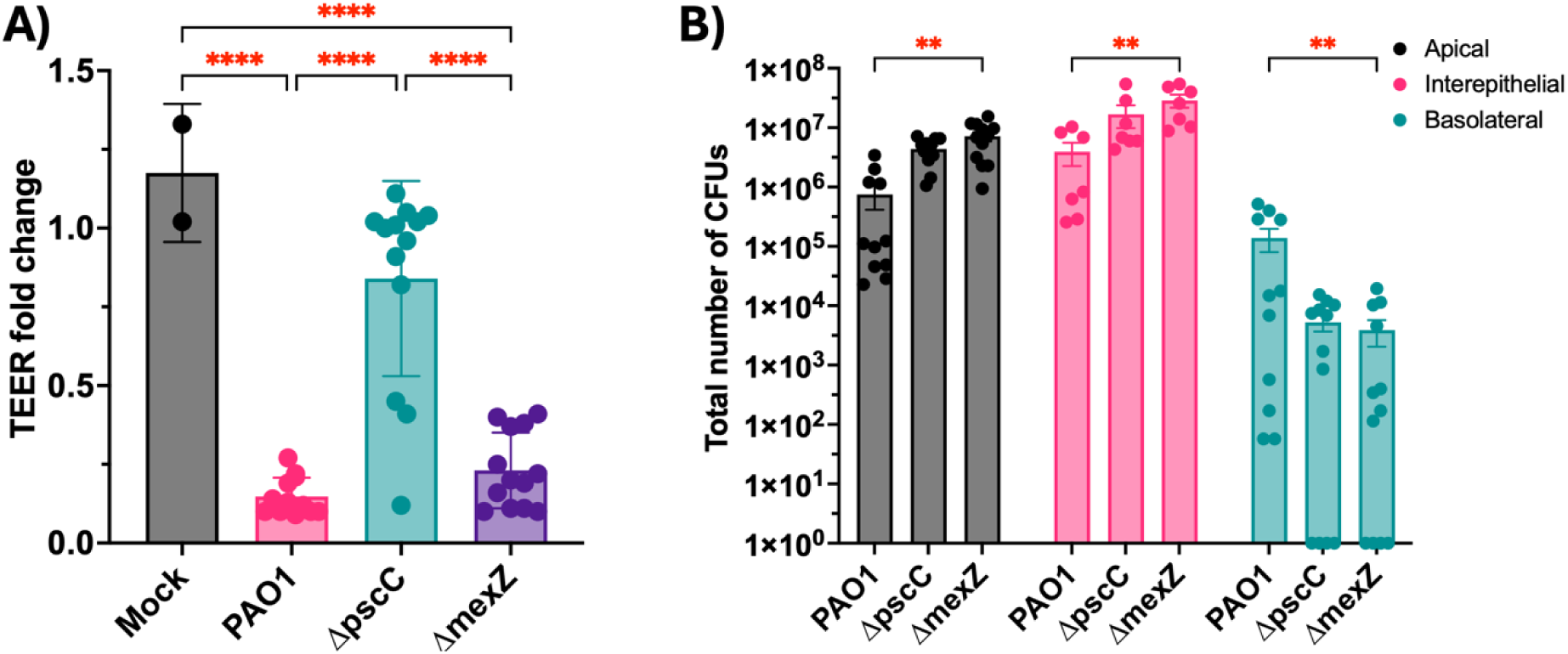
TEER measurements and CFU numbers after *in vitro* infection of lung epithelia. A) The graph depicts the fold change in TEER before compared to after infection on the y-axis; an indirect measure of degradation of tight junctions, which indicates the barrier tightness and thus “health status” of the lung epithelium. B) CFU numbers after the 16 h of incubation for each reporter strain with the basal cell immortalized-nonsmoker 1 cell line (BCi-NS1.1)-derived lung epithelia. The Y-axis is log-transformed (log(CFU+1)) for simplicity in visualization, but the two-way Analysis of Variance test (ANOVA) was carried out on untransformed data. Each dot represents one biological replica (one Transwell) across the two infection experiments. Biological significance levels were marked by highlighting asterisks in red, and is considered sufficiently important in this study, indicating a difference in TEER fold change or CFU numbers by ≥15 %. Statistical non-significance is not shown.

### 2.2 Dual-reporter strains exhibit fluorescence, have no growth defects and fluorescence decays over time

Dual-reporter strains were constructed to quantify transcription activity from a ribosomal promoter as a proxy for bacterial growth. These reporter systems were introduced in PAO1, *ΔpscC* and *ΔmexZ* to investigate bacterial growth activity in reference to tissue localization. All included *P. aeruginosa* strains had similar growth rates (Supplementary Figure 1A-D). Fluorescence developed in all strains harbouring an unstable green-fluorescent protein, but fluorescence was slightly higher from the PAO1-gfpmut3* strain harbouring a stable gfp variant (Figure 2A). Fluorescence from *Escherichia coli* strains was in general lower than what was determined for *P. aeruginosa* strains (Figure 2D), and all other strains had lower fluorescence than PAO1-superfolderGFP (sfGFP; Figure 2). This was however expected, as sfGFP is not driven by the *rrnB* promoter but by a constitutive *tac* promoter, and because sfGFP has a 1.67x higher quantum yield than the gfpmut3 variant. The PAO1/*ΔpscC/ΔmexZ*-gfpAGA strains exhibited near-identical decay rates and their half-lives were determined as exemplified by the experimentally determined Δ*pscC* decay function shown in Figure 2E: ∼60 min.

**Figure 2:**
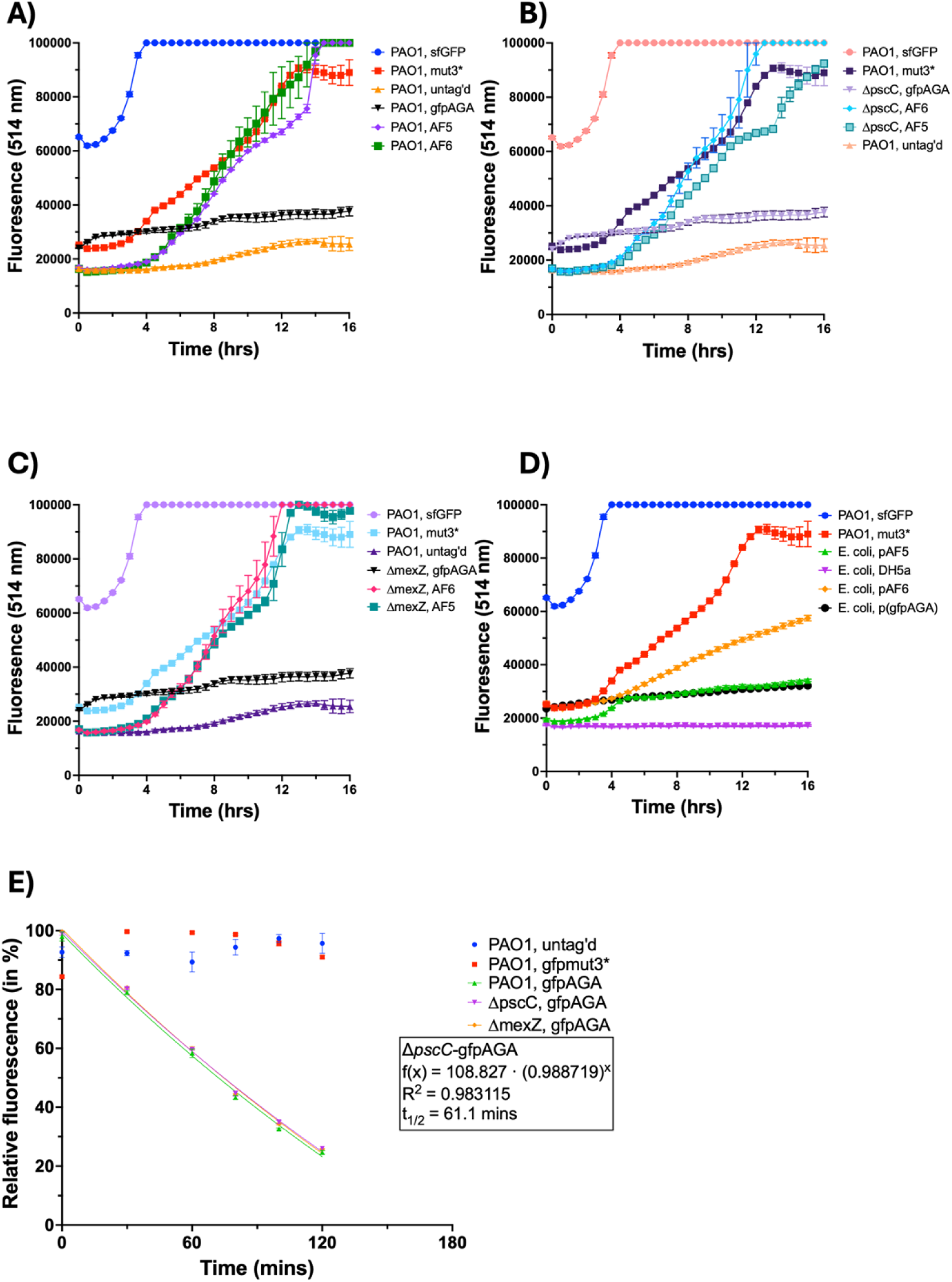
Fluorescence curves during bacterial growth and fluorescence decay after growth arrest. The Figure shows growth curves for various strains utilized in this study (panels A-D), their initial fluorescence gain at the beginning of incubation in shaking cultures (E) and the relative decay of gfpmut3*[AGA] after shift to minimal media. At around 8 h, the expected-non-fluorescent WT PAO1 began to achieve fluorescence, which is thought to be caused by production of pyoverdine and other naturally fluorescent *P. aeruginosa* proteins.

### 2.3 Microscopy reveals strain-specific distributions of bacteria across epithelial micro-niches

Confocal microscopy observations showed that bacterial populations were localized in four distinct micro-niches (Supplementary Figure 2): 1) “apical” characterized by a lack of epithelium breach and coherent actin signal, 2) “interepithelial” characterized by none-to-little tissue destruction and bacteria located between host cells, i.e., between intact “actin shells” visualized by a phalloidin-Alexa fluorophore conjugate, 3) “intracellular” characterized by bacteria located inside host cells, and in 4) “breach sites” characterized by general and widespread tissue destruction and bacteria throughout the site, and with dead host cells.

Comparing the localizations of the three *P. aeruginosa* dual-reporter strains, it was evident that the reference strain PAO1 frequently formed aggregates on the apical surface of the tissue cultures (shown in Figure 3A) and in sites associated with epithelial breaches. Populations of *ΔpscC* were frequently observed in either apical or intracellular sites (Figure 2B). Finally, microscopy investigations confirmed previous findings (16) that *ΔmexZ* bacteria were most frequently found in the interepithelial space (Figure 3C). See Supplementary Figure 2 for additional micrograph replicates and for detailed visualizations of intracellular- or interepithelial aggregates.

**Figure 3:**
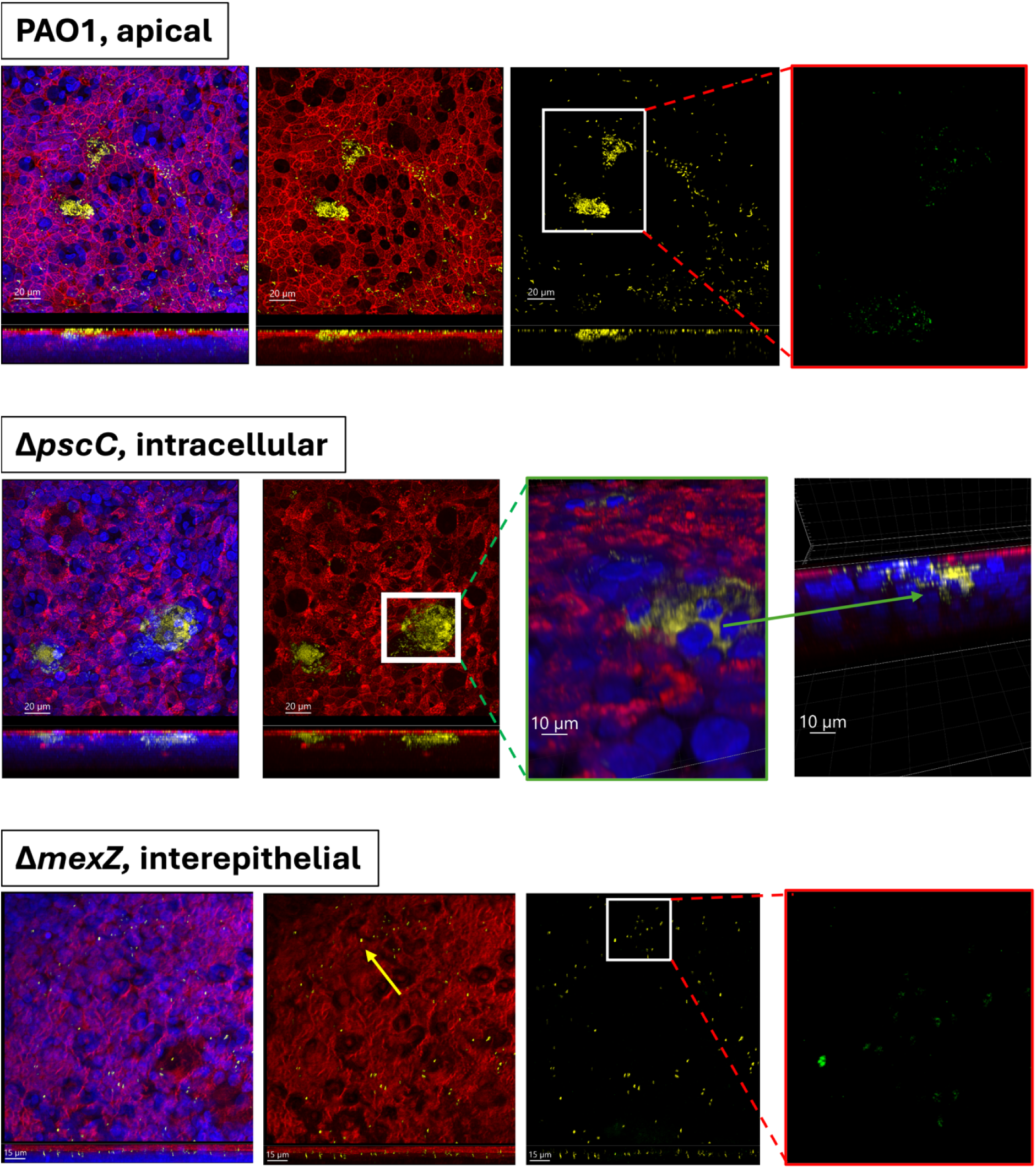
Micrographs from each dual-reporter strain as representative examples. Micrographs from confocal microscopy of A) PAO1, an example of an apical aggregate site. B) *ΔpscC*, an example of an intracellular aggregate site – note that the third and fourth images are 3D projections indicating an intracellular aggregate site, with the fourth panel being a cross-sectional 3D view of bacteria surrounding an epithelial nucleus, pointed to by a green arrow. C) example of a *ΔmexZ* interepithelial aggregate site. Note that the second panel is a cross-section in a sectional view, showing singlet bacteria spread in between F-actin “shells” (in red) of host cells, but not inside host cells. An example of such a singlet bacteria is pointed to by a yellow arrow. BCi-NS1.1 cells’ nuclei are shown in blue, actin filaments in red, Kusabira-Orange κ (mKOκ) in yellow (constitutive fluorophore expression) and the unstable green fluorescent proteins (uGFPs) in green (unstable fluorophore expression). For micrographs of PAO1 and Δ*mexZ*: Going from left to right panel: 1) BCi-NS1.1 cells (F-actin + nuclei) and bacteria are shown. 2) 1) BCi-NS1.1 cells (F-actin; no nuclei) and bacteria are shown 3) only bacteria (mKOκ and uGFP) are shown. 4) only uGFP-expressing bacteria are shown in a zoom-in of the indicated image section. The three first images of each example are shown as top-view of a total image projection of the ZStack, and in the bottom of the image, a side-view of the total image projection. See Supplementary Figure 2 for additional micrograph replicates and for more detailed visualizations of intracellular- or interepithelial aggregates.

All three strains were able to colonize each of the four micro-niches, although some were less frequently, or rarely, observed: intracellular PAO1, epithelial breach site- and interepithelial aggregates of Δ*pscC*, and apical- and intracellular aggregates of Δ*mexZ*. This distribution may however partly reflect an experimental design limitation of analysing only a single time point post-infection, as relative frequencies of micro-niche localizations might differ at earlier or later stages. It is however important to keep in mind that infection does not occur synchronously in all micro-niches: even at one time point, bacterial populations at different developmental stages coexist.

### 2.4 Quantification of growth activity by image analysis and comparison between strains revealed distinct niche preferences

The *P. aeruginosa* strains – PAO1, *ΔpscC* and *ΔmexZ* – were all tagged with stable, constitutively expressed mKOκ to visualize the localization of the bacteria used as an internal reference for quantification of uGFP signals. The uGFP is transcribed from an rRNA promoter to indicate bacterial growth activity. To determine the bacterial growth activities in the four different micro-niches, and to investigate whether the pathoadaptive mutations significantly impact growth of the three strains, a two-way ANOVA test of growth activity was performed (Figure 4).

**Figure 4:**
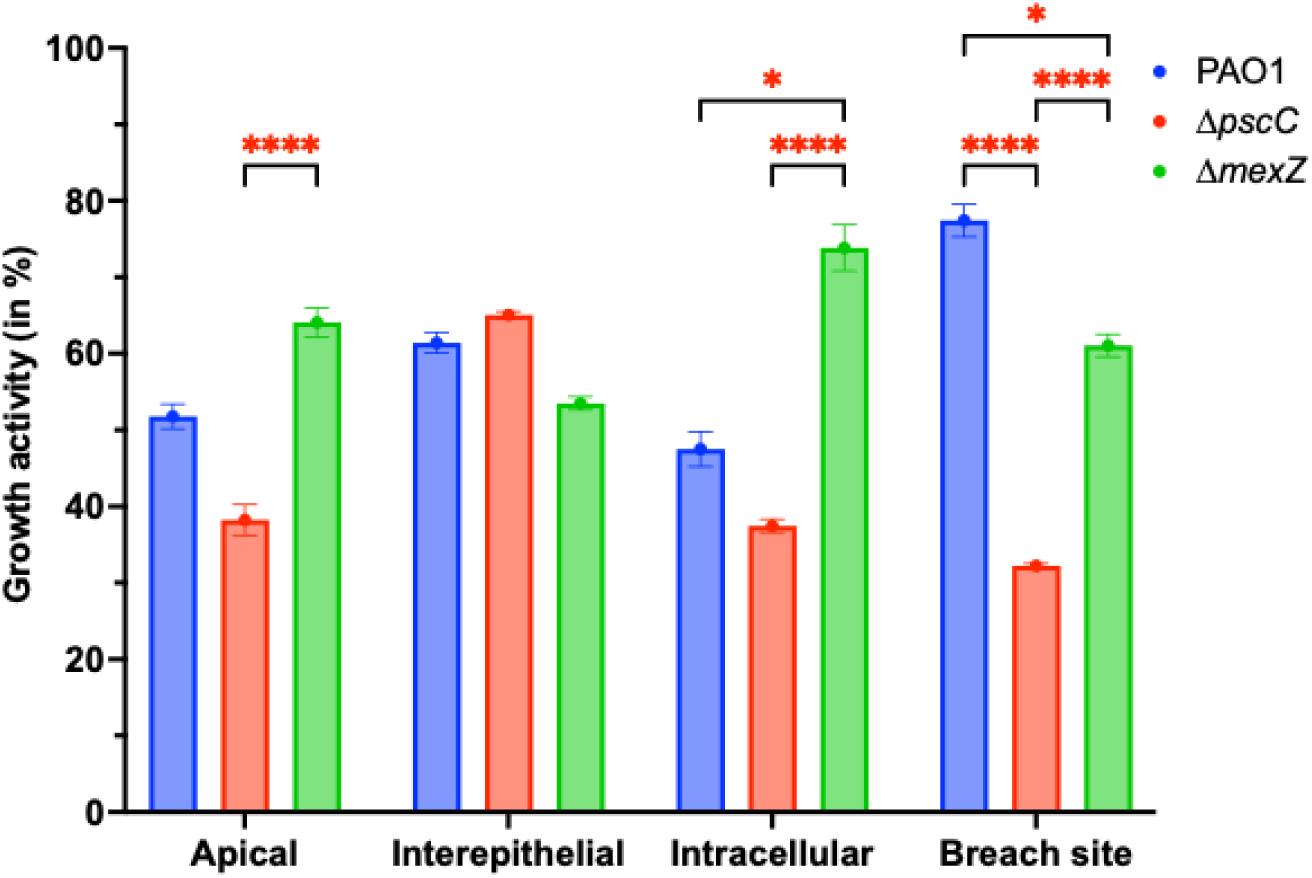
Average growth activity in each micro-niche and of each bacterial strain. The Figure depicts the average growth activity – indicated by uGFP expression relative to the top 10 % most active bacteria observed, across all recorded actively growing bacteria – divided into micro-niches and separated by bacterial strain. Only statistical comparisons *within* micro-niches and across strains are shown here. PAO1 is shown in blue, *ΔpscC* is shown in red and *ΔmexZ* is shown in green. The data is visualized as mean ± SEM. Biological significance levels were marked by highlighting the asterisks in red, and is what is considered sufficiently important in this study, indicating a difference in growth activity by at least 15 %-points. Other statistically significant (but not biologically significant) results are shown in Supplementary Figure 5. Non-significant comparisons are not shown. The number of observations in each group varied and was as follows: PAO1-Apical (n = 90). *ΔpscC-*Apical (n = 226). *ΔmexZ-*Apical (n = 91). PAO1-Interepithelial (n = 100). *ΔpscC-*Interepithelial (n = 734). *ΔmexZ-*Interepithelial (n = 170). PAO1-Intracellular (n = 11). *ΔpscC*-Intracellular (n = 59). *ΔmexZ-*Intracellular (n = 96). PAO1-Breach site (n = 135). Δ*pscC*-Breach site (n = 191). Δ*mexZ*-Breach site (n = 116). An observation is defined as a 3D object from a ZStack, regardless of how many bacteria were present in the 3D object.

PAO1 and *ΔmexZ* generally showed higher growth activities than *ΔpscC* in all micro-niches, except for interepithelial sites (Figure 4, Supplementary Figure 5). At intracellular sites *ΔmexZ* bacteria showed higher growth activity than PAO1 and *ΔpscC –* and higher activity than *ΔpscC* at apical sites. Conversely, PAO1 showed higher activity than either mutant strain at breach sites (Figure 4).

Investigations of the growth activity levels for each micro-niche and strain showed that activity levels differed significantly within each strain (Figure 5). As such, the growth activity of each strain and micro-niche could be ranked, forming hierarchies of growth activity, indicating preferred micro-niches listed in Table 1. Seemingly, pathoadaptive mutations do not only change the colonization patterns, but also the preference for sites in which they respectively exhibit highest growth activity (see Table 1).

**Figure 5:**
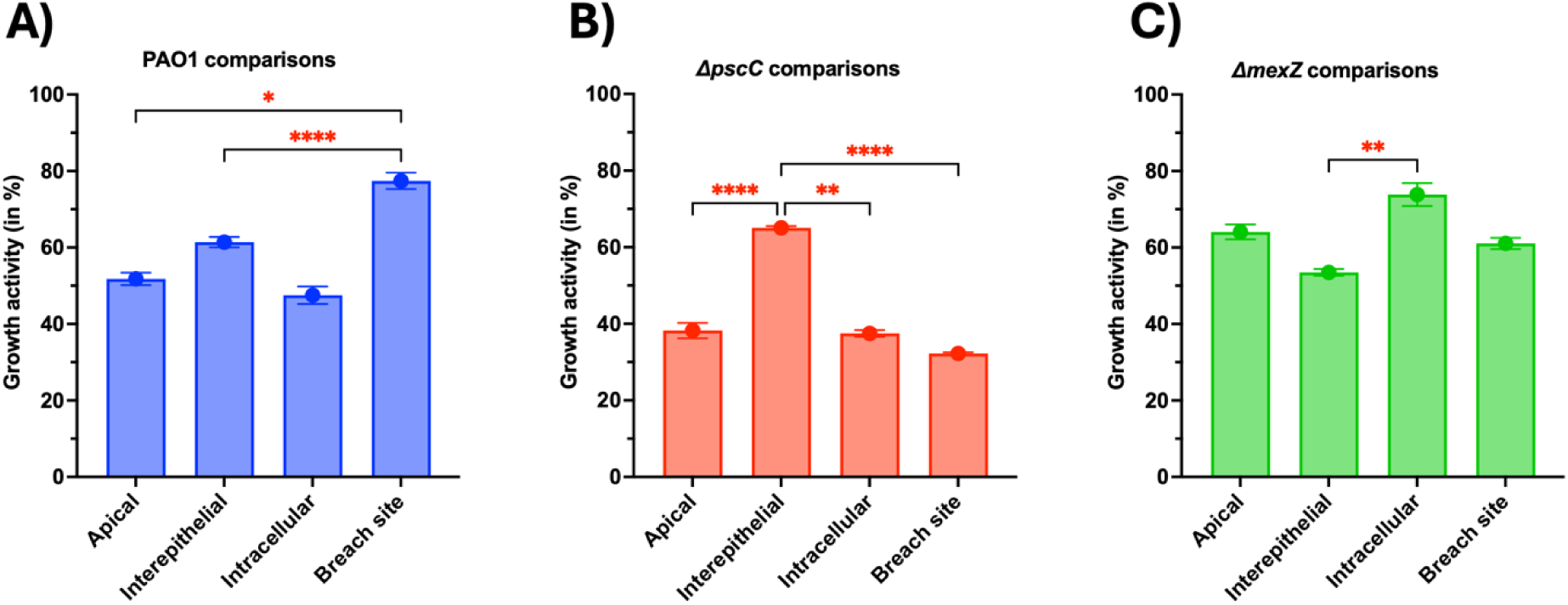
Average growth activity in each micro-niche and separated into panels depicting each respective bacterial strain. The Figure depicts growth activity, indicated by uGFP reporting, of the three strains in each of the four micro-niches. The data is visualized as mean ± SEM. Note that these three Figures were analysed together in a multiple-comparison two-way ANOVA, with a Šídák’s multiple comparisons test (shown in Figure 4), but shown individually here for the sake of simplicity. Biological significance levels were marked by highlighting the asterisks in red, and is what is considered sufficiently important in this study, indicating a difference in growth activity by at least 15 %-points. Other statistically significant (but not biologically significant) results are shown in Supplementary Figure 5. Statistically non-significant comparisons are not shown.

**Table 1.**
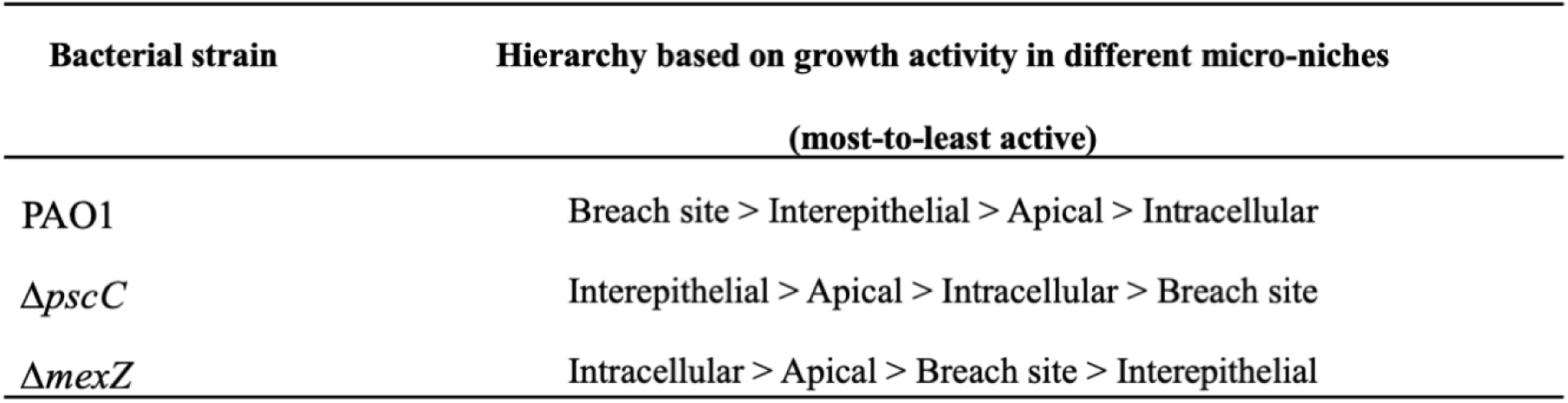
Hierarchies of growth activity for PAO1/Δ*pscC*/Δ*mexZ* in different micro-niches, ranked from most to least active site (interpreted from Supplementary Figure 5)

| Bacterial strain | Hierarchy based on growth activity in different micro-niches<br>(most-to-least active) |
| --- | --- |
| PAO1 | Breach site > Interepithelial > Apical > Intracellular |
| ΔpscC | Interepithelial > Apical > Intracellular > Breach site |
| ΔmexZ | Intracellular > Apical > Breach site > Interepithelial |

Further investigations showed that when growth activity was compared across strains without considering micro-niches, no biologically significant differences were detected (Supplementary Figure 6). However, when growth activity was compared across micro-niches independent of strain, one consistent difference emerged: populations in interepithelial sites displayed higher growth activity than those in apical sites (Supplementary Figure 7A). Interestingly, the data suggest the possibility of two distinct intracellular *ΔmexZ* populations (Supplementary Figures 7B & -11). The dual-reporter strains in combination with imaging tools also allowed for size-stratified determinations of growth activity in bacterial aggregates, as demonstrated in Supplementary Appendix 1 & -2, Supplementary Figures 3, -7, -8, -9 and Supplementary Table 1. These size-stratified analyses confirmed the findings listed in Table 1.

## 3. Discussion

In this study, we demonstrate the utility of fluorescent bacterial dual-reporter strains as a powerful tool to gain biological and spatial insights into the infection dynamics at the microscale. Specifically, we used a dual-reporter system to monitor growth activity in *P. aeruginosa* during infection of an *in vitro* lung epithelium model. This approach offers a valuable framework for dissecting the growth states and micro-niche preferences of bacterial pathogens, particularly in the context of genetic mutations relevant to virulence and persistence. Given the clinical significance of *P. aeruginosa* in airway infections in pwCF, chronic obstructive pulmonary disease, and ventilator-associated pneumonia, improving our ability to monitor its growth activity and localization in host tissues can provide important information for basic infection research as well as for therapeutic development and improvement.

Our dual-reporter strains consisted of a constitutively expressed fluorescent protein, mKOκ, serving as an internal reference and indicator of the location of the total bacterial population, in combination with an unstable, growth activity-dependent reporter, uGFP. This design enables visualization of the bacteria and their localization in the tissue cultures, and to what degree – and precisely where – bacterial growth is occurring. These reporters thus provide a quantitative and spatially resolved view of bacterial growth activity that can be computationally analysed, enabling a detailed monitoring of the infection dynamics.

Quantitative analyses of growth activities revealed distinct strain-specific growth preferences across tissue sites. PAO1 populations showed the highest growth activity at epithelial breach sites, whereas the Δ*mexZ* strain exhibited particularly elevated activity as intracellular populations. The Δ*pscC* strain showed a pattern similar to Δ*mexZ* but with the highest growth activity in interepithelial locations. Conversely, different hierarchies of growth activity across micro-niches may reflect strain-specific micro-niche preferences and distinct strategies of host colonization and virulence. It is plausible that the lowest site of activity may reflect initial attachment sites, whereas the highest may represent the point of the infection process at which most host-cell degradation is ongoing, as nutrients consequently would be released to fuel bacterial growth activity. Alternatively, it might instead reflect the availability of preferred nutrients needed for growth, for which the strains’ preferences may have changed due to pleiotropic effects of their respective mutations. Thirdly, it could be explained by the strains’ abilities to *obtain* specific nutrients in the respective preferred micro-niches, rather than the availability of them, and none of these hypotheses are mutually exclusive. Last but not least, it is important to keep in mind that the patho-adaptive mutations accumulate in an airway environment with antibiotics present most of the time. Under such conditions fast growth is not necessarily desirable for the bacteria.

*P. aeruginosa* may follow a progressive infection route, attaching to the apical surface, building up consortia of bacteria, breaching the epithelial barrier, likely through toxin-mediated damage, then disseminating via interepithelial spaces and, transiently, be located intracellularly in host cells as a part of epithelial tissue destruction, as deemed by the low frequency of observed intracellular aggregates for PAO1. Introduction of the pathoadaptive mutations *ΔpscC* or *ΔmexZ* appears to confer staggered or modified versions of this infection program. Phenotypically, Δ*pscC* bacteria were often observed intracellularly, consistent with previous reports (21,30,31). However, their growth activity at intracellular sites was relatively low (∼37%), whereas it peaked in interepithelial populations (∼65 %), suggesting that, rather than tissue invasion involving simultaneous interepithelial- and intracellular colonization, the *ΔpscC* mutant is internalized and remains in a low-activity state, until host cell death and bacterial release (32). After host cell lysis, released nutrients may cause a growth activity peak during interepithelial spread, explaining the differences observed in growth activity in these two micro-niches for *ΔpscC*.

In contrast, the *ΔmexZ* mutant strategy – as judged from the high frequency of interepithelial bacterial populations observed by microscopy and its low growth activity at these sites – seems to be initial colonization within interepithelial sites, confirming prior reports (16), which may then putatively be followed by epithelial barrier breaching and subsequent dissemination. The dissemination may involve intracellular invasion, where the highest growth activity was observed for the *ΔmexZ* mutant. The hypothesis that different low-activity colonization sites may represent initial steps in the infection process, rather than random initial colonization sites, could be tested by time-lapse monitoring from bacterial attachment to epithelial breach and dissemination. The *in vitro* tissue model applied in this study comprises fully differentiated airway cells with functional cilia, and all three bacterial strains have intact motility apparatus, further lowering the chance that the different micro-niches with lowest growth activity represent coincidental attachment sites, and rather, represent different initiation steps of the infection process.

In this study, intracellular invasion events by PAO1 were rarely observed, which is in contrast to reports by Swart *et al*. (32), perhaps due to differences in the applied cell lines, the single time-point post-infection and a relatively lower inoculum to initiate infection in this study, or likely because PAO1’s virulence level results in only a short-lived intracellular stage that is difficult to capture at a single time point. Swart *et al.* also observed frequent intracellular invasions by PAO1 in primary cells from various donors, which may indicate a limitation of the BCi-NS1.1 cell line. However, in patient samples, intracellular *P. aeruginosa* has been reported only at low frequency (33). Also, in contrast to Swart *et al.*’s study, we did not observe large intracellular populations of *ΔpscC* relative to PAO1 or *ΔmexZ,* except for one out of three ZStacks containing intracellular *ΔpscC* (Supplementary Figure 4C & Supplementary Table 2).

Interestingly, when investigating the growth activity of *ΔmexZ* intracellular bacteria, it seems as if two populations are present: one with high growth activity, another with low growth activity (Supplementary Figures 7B & 9). This has previously been reported for *P. aeruginosa*, and was suggested to be due to the presence of bacterial subpopulations in different intracellular locations (30,34), which could be investigated in future studies. The phenomenon has also been reported for *Salmonella enterica* serovar Typhimurium, using unstable reporter systems in infected epithelial cells (35). Why only the *ΔmexZ* strain exhibited polarized growth activity in intracellular subpopulations remains an open question and a potential avenue for future investigations.

The inactivation of *mexZ* leads to derepression of the *mexXY* efflux pump which is known to cause widespread pleiotropic effects (16). Previous work has shown that altering efflux pump expression can profoundly reshape the exo-metabolome (36). Such changes could modulate host-pathogen interactions during intracellular colonization, driving heterogeneous growth responses. In addition, *ΔmexZ* strains have been reported to display altered quorum-sensing-regulated pathways that contribute to their colonization phenotype in epithelial tissues (16). Within the confined environment of the host cells’ cytosols or perinuclear vacuoles (34), quorum-sensing signals may accumulate to high local concentrations. This could amplify differential activation of quorum-sensing pathways, generating distinct high- and low-growth-activity subpopulations. Other, not mutually exclusive, explanations may also apply. For instance, variability in intracellular niches, such as exposure to cytosolic versus vacuolar environments and their available nutrients, or to differing host stressors, might disproportionately impact the *ΔmexZ* mutant in comparison to PAO1 or the *ΔpscC* mutant.

Looking forward, the versatility of this dual-reporter system can prove useful in several applications. Unstable fluorescence reporters can be utilized to quantify other pathways of interest; biofilm formation (8–11), toxin expression (4,5,13), stress responses (37), two-component systems (38), motility expression (12,39), quorum sensing (14,40,41), antibiotic resistance (42), alginate production and beyond (6,7,43–45). In the future, combining such reporting with live-cell imaging will enable dynamic, real-time visualization of bacterial activity or gene expression across multiple pathways simultaneously. Such multiplexing could deepen our understanding of the temporal coordination of bacterial virulence and survival mechanisms, particularly during treatment with therapeutic compounds of interest.

Finally, growth activity reporting may hold particular relevance for development and investigations of antimicrobials. By revealing whether a potentially therapeutic compound selectively kills active or dormant bacterial populations, or both - or modifies a bacteria’s growth activities - could help elucidate drug efficacy and suggest treatment strategies, particularly against persistent or biofilm-associated infections. While constitutive reporters can reveal bacterial localization and consortia development, only unstable, activity-based reporters provide insight into the dynamic physiological states and gene expression underlying the infection. The approach presented here thus provides a valuable complement to classical imaging and quantification methods, offering a more dynamic and mechanistic view of host-pathogen interactions.

## 4. Materials & Methods

### 4.1 Bacterial strains

In this study, *P. aeruginosa* strains PAO1 (a standard laboratory reference strain (46)), PAO1-Δ*mexZ* (16) and PAO1-Δ*pscC* (47) were applied as background strains for genetic transformation procedures with the dual-reporter system. All bacterial strains utilized in this study and their GFP/mKOκ variants, if applicable, are listed in Supplementary Table 3.

For growth- and fluorescence curves, a PAO1sfGFP strain was applied. The strain was constructed from a sfGFP gene fragment which was PCR-amplified from the plasmid pSEVA628 using primers containing SacI and KpnI sites. The amplified fragment was cloned into the mini-Tn7 delivery plasmid pJM220 (48) using the same restriction sites. Successful ligation was obtained from a 3:1 insert-to-vector ratio along with TSAP (Thermo Sensitive Alkaline Phosphatase) treatment, to avoid plasmid recircularization. Plasmid was transformed into chemically competent *E. coli* DH5ɑ, and positive transformant colonies were selected on lysogeny broth (LB) agar plates containing 100 µg/ml ampicillin.

### 4.2 Cloning Procedures

Dual-reporter strains, termed -AF5 or -AF6, were constructed by genomic integration of genes encoding two fluorescent proteins. A pUC-derived mini-Tn7 plasmid was constructed, which encoded a uGFP (gfpmut3*-AGA for AF5 (23,25,49–52) or sfGFP-AGA for AF6 (53)) and stable mKOκ fluorescent protein, both derived from pJM220. The plasmids were then named pAF5/pAF6. Synthetic uGFP gene fragments were purchased from, and synthesised by, Integrated DNA Technologies (IDT, Europe), and inserted into the plasmid via restriction enzymes NsiI and HindIII. uGFP expression is controlled by the *rrnB* promoter, P1; a growth-rate regulated *E. coli* ribosomal promoter (23,24). This leads to expression of the uGFP during bacterial growth, and uGFP is degraded over time due to its C-terminal peptide sequence -AGA (11,23,45,49–52), ensuring differential signal intensity based on recent promoter activity. In contrast, the mKOκ expression is controlled by a constitutive *tac* promoter, ensuring constant reporting on bacterial location, regardless of growth activity.

The genetic constructs were designed to contain NsiI and HindIII restriction sites, enabling easy cloning into the mini-Tn7 transposon plasmid (25). Chemically competent *E. coli* DH5ɑ harbouring the pAF5 or pAF6 plasmids were genetically transformed and positive transformant colonies were selected on lysogeny broth (LB) plates containing 10 µg/ml gentamicin.

### 4.3 Construction of Reporter Strains

The PAO1sfGFP and dual-reporter strains were constructed by tetra-parental mating using the Tn7 transposition system with gentamicin resistance selection (30 µg/ml) on *Pseudomonas* isolation agar, as described previously (25,54,55). This ensures gene insertion in one direction, at a unique and neutral chromosomal site, downstream of the glucosamine synthetase gene, *glmS* (25). Importantly, the system is *not* plasmid-based but is integrated in the genome as a single copy, meaning there are no quantification biases introduced from potential plasmid copy numbers. The new reporter strains were named PAO1-AF5 or -AF6, PAO1-Δ*pscC*-AF5 or - AF6 and PAO1-Δ*mexZ*-AF5 or -AF6, and referred to simply as “PAO1”, “Δ*pscC”* or “Δ*mexZ*” for the remainder of this article. Gfp-AGA expression in PAO1 has been described in detail previously (23,50).

### 4.4 Culturing of Human Lung Epithelial Tissue

BCi-NS1.1, an immortalized human airway basal cell line, was cultured statically at two different days – leading to two experimental replicas – in fifteen and twenty-three Transwells of 24-well Transwell plates (0.4 µm pore size, Corning), respectively, and cultured as described elsewhere (16). The Transwells were infected with the constructed reporter strains of PAO1, Δ*pscC* and *ΔmexZ* after culturing of the human BCi cells for 28 days at air-liquid interface. The first experimental replicate contained five biological replicas of each reporter strain, whereas the second experiment comprised seven PAO1 replicas, and eight of the *ΔpscC* and *ΔmexZ* strains, each. An un-infected control well (“Mock”) was included in both experiments, which received PBS instead of bacteria, at the initiation of infection.

### 4.5 Infection Experiments

TEER was measured 3 h prior to infection start, simultaneously with basolateral media replenishment. The reported fold-change in TEER was calculated as TEER_post-infection_/TEER_t0_ and gives an indication of the degradation of tight junctions, as high TEER values indicate strong tight junctions between cells. Afterwards, 10 µl of 10^4^ CFU/ml bacterial suspensions, diluted in sterile phosphate-buffered saline (PBS), was placed in the centre of each Transwell, followed by incubation at 37 °C and 5 % CO_2_ in a humidified incubator. Sixteen hours later, 200 µl PBS was added to the apical Transwell chamber, TEER was measured again, the apical PBS was collected (containing the non-attached, planktonic, bacterial population), and two Transwells of each strain were fixed by addition of 4 % paraformaldehyde (aq.). The remaining Transwells of each strain were added 200 µl of 0.02 % Triton-X (in PBS) and incubated for 20 min at 4 °C followed by manual cell homogenization. This compartment was sampled to quantify attached *P. aeruginosa* numbers. Basolateral media was collected from all Transwells (containing the fraction of bacterial population which successfully penetrated the epithelium and disseminated into the basolateral media).

### 4.6 Bacterial enumeration

Inocula from the static Transwell infections were diluted to a concentration of 1×10^3^ CFU/ml and six droplets of 10 µl each was plated onto LB plates, immediately after inoculation of the Transwells. These were used to verify the number of inoculating CFUs.

Post-infection, 10 µl droplets of dilution series from each Transwell (undiluted sample to 10^-6^ dilution) was plated on LB plates, dried, and incubated for 16-20 h at 37 °C prior to CFU counting. As 10 µl was plated of the undiluted sample, the limit of detection was 100 CFU/ml. Because the liquid volumes of apical- and interepithelial compartments were 200 µl PBS each, a minimum of 20 CFUs would be detectable, and minimum 40 CFUs would be detectable in the basolateral compartment, from which the total volume was 400 µl.

### 4.7 Confocal Laser Microscopy & Fluorescence detection

uGFP was visualized by excitation at 470 nm to avoid crosstalk between fluorescent proteins, and emission was detected at 501-540 nm, whereas mKOκ fluorescence was visualized by excitation at 551 nm and its emission detected at 560-600 nm. Lung epithelial cells’ nuclei were visualized by DAPI (excitation 405 nm and emission 410-500 nm, ThermoFisher) and F-actin by Phalloidin-Alexa 647 (excitation 650 nm and emission 660-700 nm, ThermoFisher).

### 4.8 Growth curves, fluorescence curves and fluorescence decay experiments

To determine growth- and fluorescence curves, strains (PAO1-gfpmut3* (43,56), untagged PAO1 (46), PAO1-gfpmut3*-AGA (50), PAO1/*ΔpscC/ΔmexZ*-AF5, PAO1/*ΔpscC/ΔmexZ*-AF6, PAO1-sfGFP, *E. coli* strains DH5a, -pAF5, -pAF6, and -p(gfpAGA) (50) were incubated in 200 µl LB media at 37 °C for 16.5 h on a microtitre plater reader (Agilent BioTek Synergy H1), each in tri- or quadruplicates, starting at an optical density of 0.1 (measured at 600 nm; OD_600_, Supplementary Figure 1). Every 30^th^ min, OD_600_ was measured and GFP fluorescence was read by excitation at 488 nm and emission at 513 nm (optimum emission), to yield growth-and fluorescence curves.

Fluorescence decay of the respective unstable gfpAGA proteins were measured via an adapted protocol from Andersen *et al.* (43). Briefly, the cultures were diluted to an OD_600_ = 0.1 in 25 ml LB media, in a 250 ml Bluecap round-bottom culture flask, and were cultured at 200 rpm and 37 °C. At OD_600_ = 1, bacteria were pelleted and resuspended in 25 ml M9 media (without addition of a carbon source) and returned to new culturing flasks at 200 rpm and 37 °C. Fluorescence was measured every 20^th^ min and was detected by excitation at 506 nm – which is the excitation optimum for gfpmut3* – and emission at 535 nm. However, the first two measurements were made after 30 and 60 min, respectively. The fluorescence half-life was calculated as shown in Equation 1.

**Equation 1.** Determination of half-life, t_1/2_, of unstable fluorescent proteins.

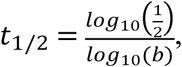

where b is the decay factor given in the experimentally determined decay function, *y = a·b^t^*

### 4.9 Image analysis

Fiji was used for image analysis and quantification of bacterial numbers and their signal intensities in ZStacks (57). Firstly, if Brightness & Contrast was adjusted, and/or if noise subtraction was necessary, it was performed with identical parameters for each of the two quantified fluorescence channels (mKOκ and uGFP), prior to image analysis of individual ZStacks. All images were quantified as 16-bit images. Not all ZStacks needed image processing prior to quantification. When noise reduction was deemed beneficial, pixels were subtracted by the Process > Math > Subtract tool, in Fiji. For 3D Object Counting, the Options were set for Volume, Integrated Density, Centroid and Mean Gray Value. Many of the detected 3D objects comprised several bacteria which were not individually distinguishable by the 3D Object Counting tool. These were defined as aggregates and is what is referred to in the article as aggregates. Thus, to count bacteria in each ZStack by image analysis, single bacteria were identified manually, and the average of 4-8 single bacterias’ volumes was applied as the volume of a single bacterium. By dividing larger 3D object’s volume with the volume of a single bacterium, the total number of bacteria per 3D object was calculated. All detected 3D objects containing uGFP signal to *any* degree was counted, regardless of signal intensity, and was used to calculate the % of active bacteria (i.e., the % of bacteria containing *some* indication of growth activity out of the total population). This was calculated as shown in Equation 2. Forty-one ZStacks were analysed (12-15 ZStacks for each strain).

**Equation 2.** Calculating the % of the total bacterial population which had detectable, ongoing, growth activity at the time of sample fixation.

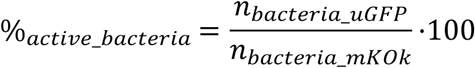

It was also possible to calculate the growth activity of each bacterium, or average growth activity of an aggregate. This was done by dividing the uGFP signal density of a 3D object with the average signal density of bacteria in the mKOκ channel, yielding a ratio of uGFP-to-mKOκ signal for each respective 3D object in a ZStack (i.e., using mKOκ as an internal reference). The average ratios of the 10 % highest uGFP-to-mKOκ ratios detected across all forty-one ZStacks were used as a reference of 100 % activity, giving a proxy for maximal growth activity which was compared across ZStacks. Note that the AF5 and AF6 dual-reporter strains were analysed separately in this manner to account for differences in quantum yield, brightness and other parameters, of their respective GFP variants.

In general, the mKOκ signal had low spread across 3D objects within ZStacks. As there were differences in the number of 3D objects detected in each ZStack (a 3D object can contain one to several thousand bacteria), the weighted average of standard deviations of the mKOκ signal was 14.3 ± 0.0016 %, indicating a quite stable mKOκ fluorescence signal, applicable as an internal reference. One reason for this variance may be the presence of two copies of the bacterial genome before cellular division at replication, particularly in sites with high growth activity.

Twelve to fifteen aggregate-containing ZStacks were quantified for each respective *P. aeruginosa* reporter strain. Of the total forty-one ZStacks, at least three replicas of each reporter strain were found for each of the micro-niches, with the exceptions that only one “breach site” was found for Δ*pscC* and only one “intracellular site” ZStack was found for wild-type (WT) PAO1.

The number of bacteria per ZStack ranged from 85 to 69,434 bacteria (average: 4,917 ± 1,789 bacteria/ZStack ± SEM). Conversely, the weighted average volume of a single bacterium was 1.16 µm^3^ (ranging from 0.72 to 1.73 µm^3^). To generate a general overview of the collective data, the average GFP activity in each ZStack was computed along with the spread (indicated by standard deviation), and the relative distribution of bacteria found as singlets (1-2 bacteria), small aggregates (2-30 bacteria/3D object), medium-sized aggregates (30-150 bacteria/3D object) or large aggregates (more than 150 bacteria/3D object). See Supplementary Table 2 for an overview of the data.

### 4.10 Statistics & Software

Statistical analyses of data from image quantification were performed in Prism 10. In Figure 1A and Supplementary Figure 6, one-way ANOVAs with Tukey’s multiple comparisons tests, were carried out. For Supplementary Figures 9 and -10, one-way ANOVA with Dunn’s multiple comparison tests were carried out. For Figures 1B, -4, -5 and Supplementary Figure 5, two-way ANOVAs with Šídák’s multiple comparisons tests were carried out. For Supplementary Figures 3E-H, -4, -7 and -8, a two-way ANOVA with Tukey’s multiple comparisons test was carried out. The exponential functions of the fluorescence decay data (Figure 2) were modelled in Prism 10. Lastly, R was used for K-Means Clustering analyses and the Principal Component Analysis shown in Supplementary Figure 3A-D. Statistical significance levels were defined as a p-value at or below 0.05 (*), 0.01 (**), 0.001 (***) and p<0.0001 (****). Biological significance levels were marked by highlighting the asterisks which show statistical significance in red, and is what is considered sufficiently important in this study to indicate a remarkable biological effect, meaning a difference in effect (or growth activity) of at least 15 %-points.

## Acknowledgements

The authors would like to thank Professor Ronald G. Cristel (Weil Cornell Medical College, New York, USA) for providing the BCi-NS1.1 cell line utilized in this study and Dr. Hanna Müller-Esparza for the generation of plasmid p(mKOκ). Our gratitude also goes to the staff at the Center for Biosustainability at the Technical University of Denmark.

## Supporting Information

**S1 Figure. Growth curves of various bacterial strains used in the study.**

**S2 Figure. Examples of ZStacks from each of the three *P. aeruginosa* strains utilized in the study (PAO1/Δ*pscC*/Δ*mexZ*), for each of the four observed micro-niches.**

**S3 Figure. Data analysis behind the rationale and decision-making for the chosen sizes of aggregates for statistical analyses in Appendix S1 and Supplementary Figures 4, 7B, 8 and 9.**

**S4 Figure. No differences were found between the fraction of the bacterial population that are singlets, small aggregates, medium-sized aggregates or large aggregates, when comparing each micro-niche within each of the three dual-reporter strains.**

**S5 Figure. Graphs of average growth activities divided into micro-niches and separated by bacterial strain.**

**S6 Figure. Independently of the micro-niche or aggregate sizes, there are no general biologically significant differences between the growth activity of the three dual-reporter strains.**

**S7 Figure. Independently of the bacterial strain, there *are* statistically significant differences between activity levels between the micro-niches under investigation. Also, it is shown that the Δ*mexZ* may harbour two subpopulations with distinct physiological states in intracellular environments.**

**S8 Figure. Comparisons of growth activity separated by the four micro-niches *and* three dual-reporter strains, divided into aggregate sizes.**

**S9 Figure. Size-stratified analyses on growth activity of different aggregate sizes of the *ΔmexZ* mutant strain at all four micro-niches.**

**S1 Appendix. Analyses on the distribution of aggregate sizes revealed no differences between strains nor between the micro-niches.**

**S2 Appendix. Size-stratified analyses reveal strain- and micro-niche-specific growth patterns.**

**S1 Table. Overview of relative growth activity (indicated by +) of *P. aeruginosa* strains across aggregate sizes and micro-niches.**

**S2 Table. Overview of all quantified parameters from the infection experiments by image analysis separated by ZStacks.**

**S3 Table. Overview of the various bacterial strains used in this study, and their respective GFP variants**

